# Structure of the human BBSome core complex in the open conformation

**DOI:** 10.1101/845982

**Authors:** Björn U. Klink, Christos Gatsogiannis, Oliver Hofnagel, Alfred Wittinghofer, Stefan Raunser

## Abstract

The BBSome is a heterooctameric protein complex that plays a central role in primary cilia homeostasis. Its malfunction causes the severe ciliopathy Bardet-Biedl syndrome (BBS). The complex acts as a cargo adapter that recognizes signaling proteins such as GPCRs and links them to the intraflagellar transport machinery. The underlying mechanism is poorly understood. Here we present a high-resolution cryo-EM structure of a human heterohexameric core subcomplex of the BBSome. The structure reveals the architecture of the complex in atomic detail. It explains how the subunits interact with each other and how disease-causing mutations hamper this interaction. The complex adopts a conformation that is open for binding to membrane-associated GTPase Arl6 and a large positively charged patch likely strengthens the interaction with the membrane. A prominent negatively charged cleft at the center of the complex is likely involved in binding of positively charged signaling sequences of cargo proteins.

## Introduction

Ciliary research had a rocky trail to travel since the discovery of cilia in 1675 as the first known organelle by Antony Van Leeuwenhoek. Although primary cilia have long been thought to be only minor players in the cellular opera ^1^, they perform key functions as cellular antennae. Primary cilia contain a plethora of crucial signaling proteins with important sensory and regulatory functions ^2, 3^. Since cilia do not contain a protein synthesis machinery, a key question of ciliary research is how proteins are transported to, from and within the cilium. The BBSome is a cargo adaptor that recognizes a diverse set of membrane-bound ciliary proteins. It binds with its cargo to the intraflagellar transport (IFT) complex, a large heterooligomeric protein complex that is transported along ciliary microtubules by the molecular motors dynein and kinesin ^4, 5^. Interestingly, the BBSome is also involved in the assembly and stabilization of the IFT complex ^6, 7^, with varying impact on IFT stability in different organisms ^6, 8–10^.

It has been proposed that the BBSome is assembled in a sequential order starting from a complex of BBS7, BBS chaperonins and the CCT/TRiC complex, which acts as a scaffold for further BBSome subunits to be added ^11^. However, this route cannot hold true in *Drosophila*, where the BBSome components BBS2 and BBS7 are missing. The lack of these domains in *Drosophila* raises in general the question whether these subunits are important for central BBsome functions such as cargo and membrane binding or if they are mostly involved in cilia-specific processes that require the interaction with the IFT complex. In line with this, *Drosophila* and other organisms with a small number of ciliated cells lack the IFT proteins IFT25 and IFT27 ^12^, which are considered to be the anchor points for the BBSome on the IFT complex ^13–15^.

BBS1 emerged to be the most important BBSome subunit for cargo recognition, with several described interactions with ciliary cargo proteins ^16–22^. The BBSome is recruited to membranes by the small GTPase Arl6 ^16^, which binds to the N-terminal β-propeller domain of BBS1. The crystal structures of this domain in complex with Arl6 ^23^, as well as the β-propeller of BBS9 ^24^, provide the only currently available high-resolution structural information on BBSome subdomains. While this manuscript was in preparation, the medium-resolution structure of a BBSome complex purified from bovine retina was reported ^25^. The structure revealed that BBS2 and BBS7 form a top lobe that blocks the Arl6 interaction site on BBS1. The complex, however, has been purified by affinity chromatography using Arl6 as bait. This suggests that the conformation of the BBsome in its apo state differs from that of the complex bound to Arl6.

Although the cryo-EM structure at 4.9 Å revealed the overall domain architecture of the complex, an atomic model could not be accurately built due to the limited resolution. However, this is needed to fully understand the interactions of the domains and the mechanism underlying cargo binding and membrane interaction.

We recently reconstituted a heterologously expressed core complex of the human BBSome, comprising the subunits BBS1, 4, 5, 8, 9, and 18 ^26^. Although this complex lacks BBS2 and BBS7, it binds with up to sub-micromolar binding affinity to cargo proteins. In addition, we found that strongly binding ciliary trafficking motifs contain stretches of aromatic and positively charged residues, many of which were located in the third intracellular loop and the C-terminal domain of ciliary GPCRs. Multiple binding epitopes on cargo proteins cooperatively interact with multiple subunits of the BBSome ^26^, which is consistent with previous reports ^16^. A detailed insight of how these motifs interact with the BBSome requires a full molecular model of the complex to evaluate the potential interaction surfaces on the side-chain level.

Here we report the structure of the human BBSome core complex at an average resolution of 3.8 Å for BBS1, 4, 8, 9, and 18 and 4.3 Å for BBS5. The high quality of the map allowed us to build an atomic model of ~80% of the complex. The structure reveals the architecture of the complex and the sophisticated intertwined arrangement of its subunits. A large positively charged region on the surface of the complex suggests how it orients on the negatively charged ciliary membrane. We found that the complex adopts an open conformation in which the Arl6 binding site would be accessible. The high-resolution structure allows us to accurately locate pathogenic patient mutations and to decipher how they perturb intra- and/or intermolecular interactions on the molecular level. We identified a negatively charged cleft in the center of the complex, which is positioned favorably for cargo interactions. This provides the first structural indication for how cargos are probably recognized by the BBSome.

## Results and Discussion

### Architecture of the BBSome core

To obtain suitable samples for high-resolution structural analysis, we chose to work on the BBSome core instead of the full complex, because the core, comprising BBS1, 4, 5, 8, 9, and 18 was considerably better soluble, more stable and homogeneous, but still bound with high affinity to Arl6 and cargo peptides (Figure 1A) ^26^.

**Figure 1:**
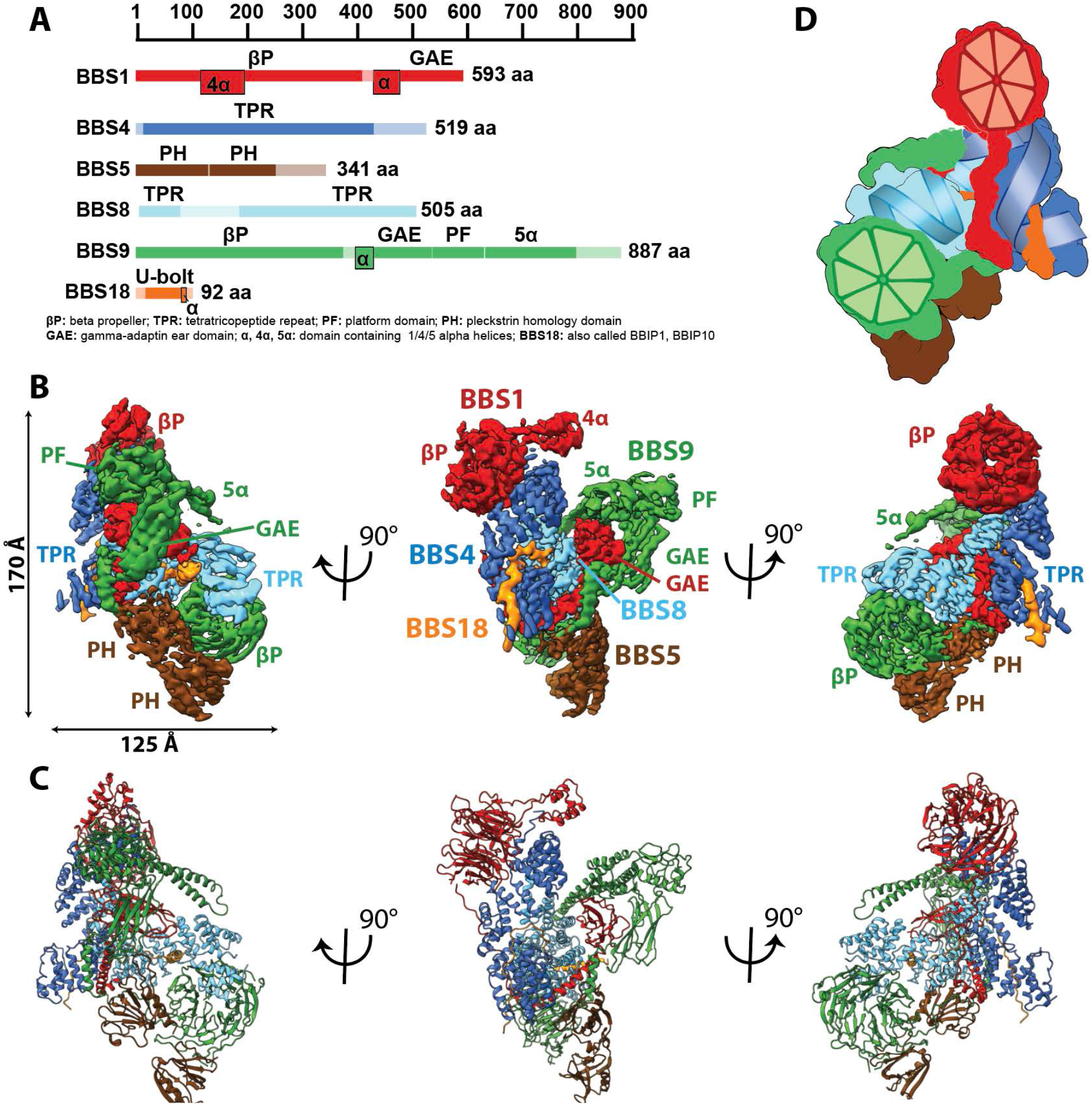
Architecture of the BBSome. **(A)**: Domain architecture of the protomers forming the BBSome core complex. The parts of the primary structure that could be assigned in the density is shown in full colors, while non-modeled regions are represented in opaque colors. **(B)**: Cryo-EM density map of the BBSome core complex in different orientations. Each protomer is colored differently and the density threshold of the individual domains were adjusted to visualize each domain at an optimal signal intensity. BBS5 is poorly visible at the signal level of the other subunits and required a reconstruction from a subset of particles, which was obtained by 3D-sorting (Figure S1). **(C)**: The final model of the core BBSome. **(D)**: Schematic representation of the complex, highlighting the two β-propeller domains of BBS1 and BBS9 as red and green segmented wheels, and the super helical arrangements of the TPR repeats of BBS4 and BBS8 as blue and cyan helices.

We determined the structure of the BBSome core complex by electron cryo microscopy (cryo-EM) and single particle analysis in SPHIRE ^27^. A reconstruction of the full data set yielded a resolution of 3.8 Å and besides BBS5 that was only sub-stoichiometrically bound (Figure S1A) all domains were well resolved with only some connecting loops and N-and C-terminal regions missing (Figure 1A, Figure S1, S3). After three-dimensional sorting (Methods, Figure S4) we also resolved BBS5 at a resolution of 4.3 Å (Figure S1) and combined its density with the reconstruction of the full data set (Figure 1B) to build an atomic model (Methods, Figure 1C, Table S1).

The BBSome core complex is arranged in multiple layers with its smallest 93 residue subunit BBS18 (also known as BBIP10, BBIP1) in its center (Figure 2). While being almost completely unfolded itself, BBS18 winds through two super-helically arranged TPR domains of the perpendicularly arranged BBS4 and BBS8 subunits, and clamps them together like a U-bolt (Figure 1, Figure 2A,B, Figure 3A). The resulting Y-shaped arrangement forms the spine of the core BBSome complex. BBS1 binds to the N-terminal end of the TPR superhelix formed by BBS4 via its N-terminal β-propeller domain (Figure 3D), wraps around BBS4 and BBS8, and binds with its C-terminal GAE domain to the TPR domain of BBS8 (Figure 2C, Figure 4A). BBS9 has a similar domain architecture as BBS1, with two additional domains at its C-terminus. The N-terminal β-propeller of BBS9 binds to the N-terminus of the TPR superhelix formed by BBS8, in analogy to the BBS1/BBS4 interaction (Figure 3C). BBS9 wraps around BBS4 and parts of BBS1, and engulfs the GAE domain of BBS1 with its C-terminal GAE, platform and α-helical domains (Figure 2D, Figure 4A). BBS5 is composed of two PH domains that both interact with the β-propeller of BBS9. One of the PH domains also interacts with BBS8 and with an unstructured loop of BBS9 (Figure 2E; Figure S1C,E).

**Figure 2:**
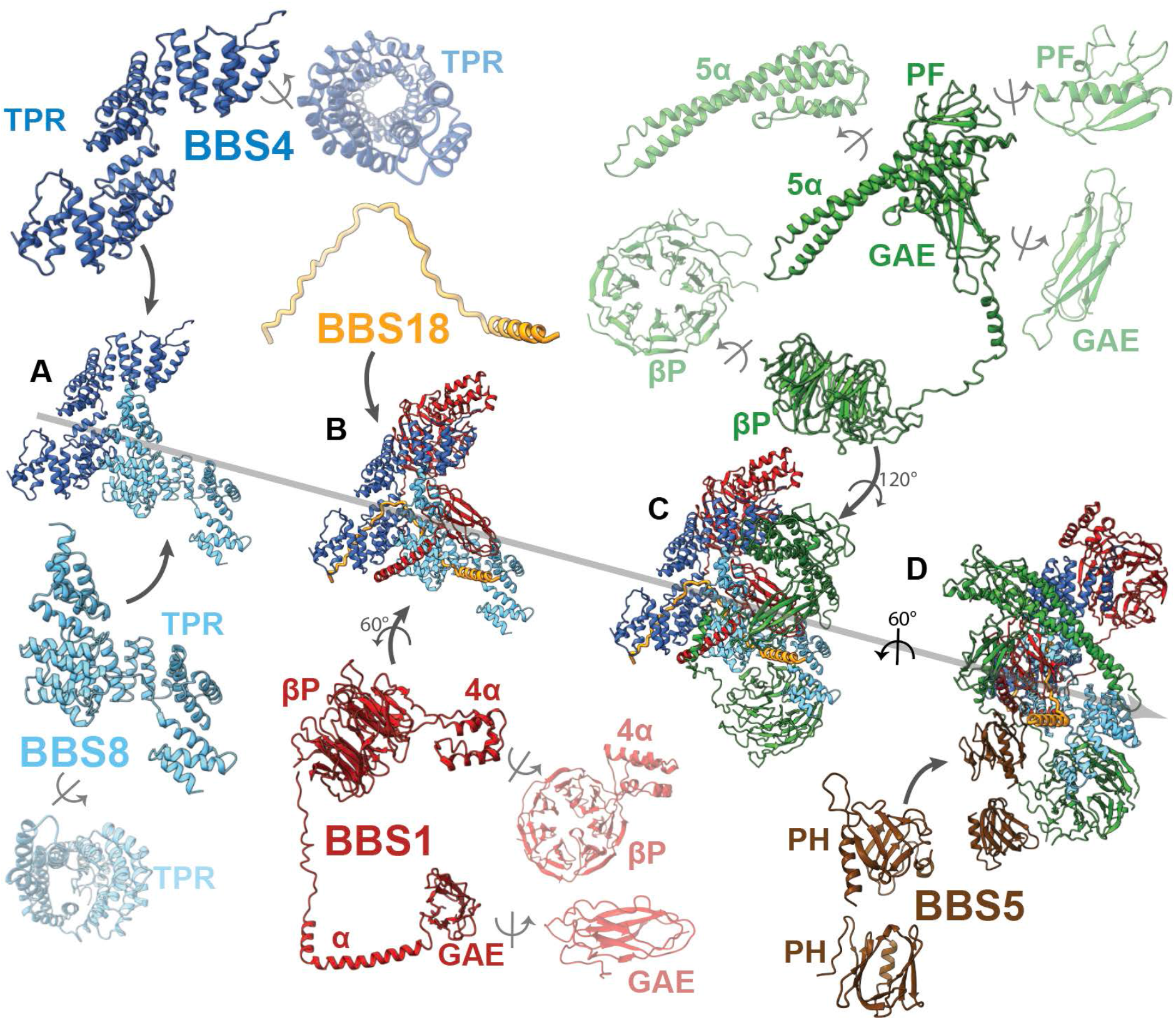
Arrangement of subunits and domains within the BBSome core. **(A-E)**: The described order of addition of subunits has been chosen for visual clarity and does not reflect the sequential assembly *in vivo*. BBS4 and BBS8 are shown in two different views to visualize the superhelical arrangement of the TPR repeats. Likewise, domains of BBS1 and BBS9 are also shown individually in two views.

**Figure 3:**
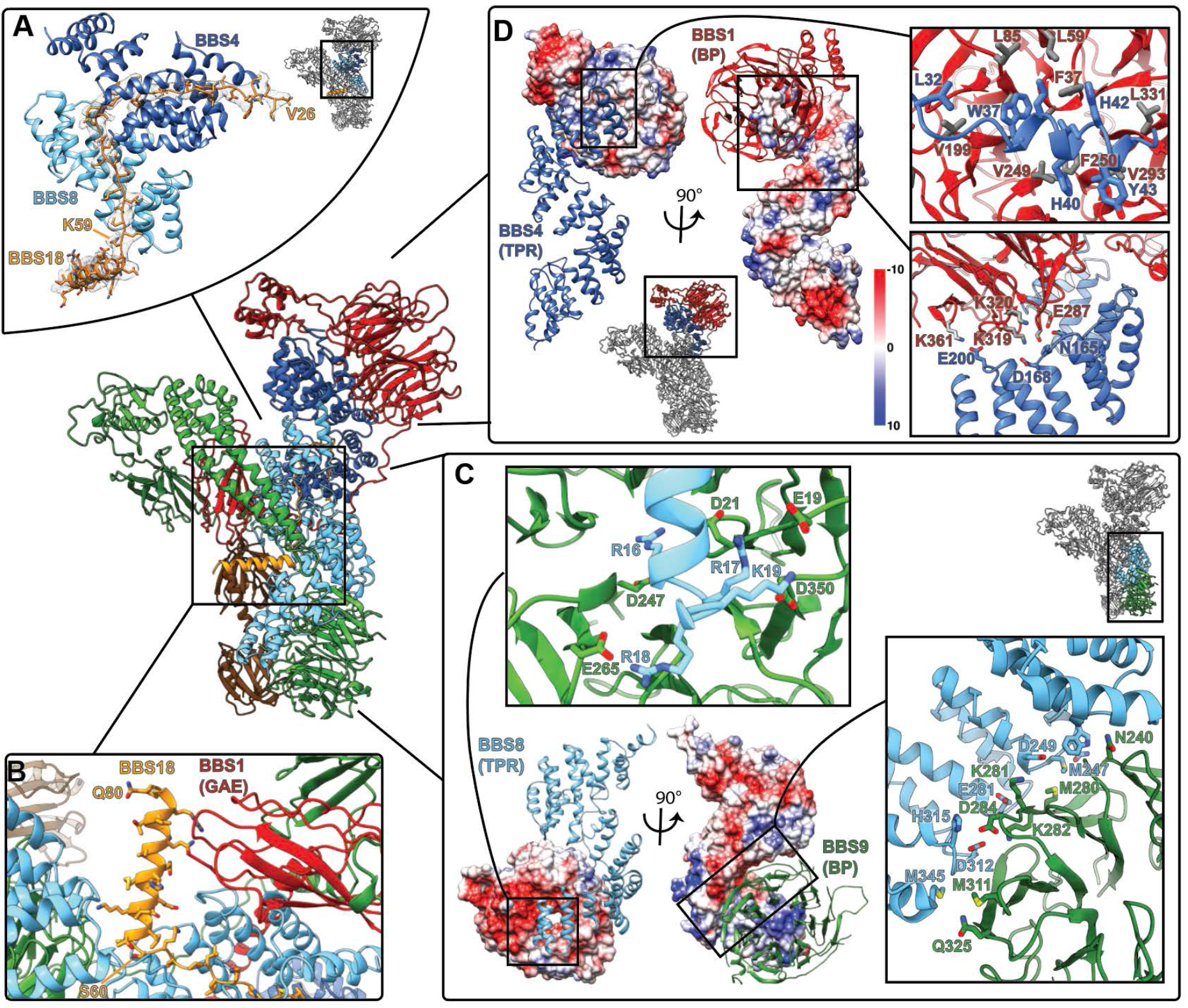
Interactions of the TPR repeat proteins BBS4 and BBS8 with BBS1 and BBS9 β-propellers and with BBS18. **(A)**: Interactions of BBS4 and BBS8 with the BBS18 U-bolt region (Val26-Lys59) stabilize the central spine of the complex. **(B)**: In contrast, the C-terminal BBS18 helix (Ser60-Gln80) only interacts weakly with the complex, and is held in place by interactions with the GAE domain of BBS1. **(C)**: Surface charge complementarity of the interaction surface of BBS8 with the β-propeller (BP) of BBS9. **(D)**: Surface charge complementarity of BBS4 with the BBS1 β-propeller.

**Figure 4:**
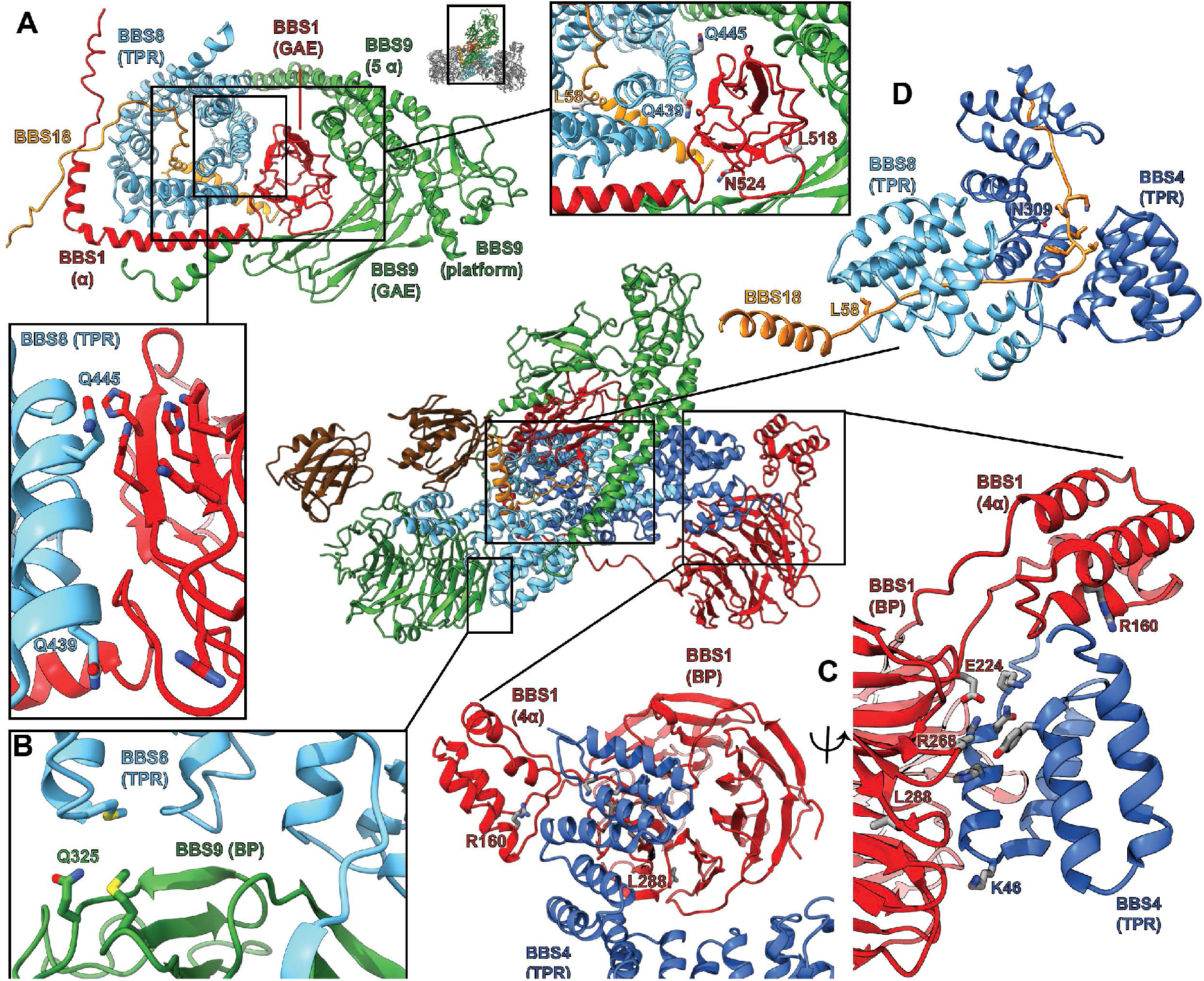
Pathogenic patient mutations within the core BBSome complex with the potential to disturb subunit interactions. **(A)**: The GAE domain of BBS1 binds to BBS8 and is embraced by the C-terminus of BBS9. Pathogenic patient mutations Q439H and Q445K on BBS8 and L518P and N524Δ on BBS1 disturb this interface. **(B)**: The pathogenic mutation Q325R in BBS9 is located at the interface of the BBS9 β-propeller with BBS8. **(C)**: Several pathogenic patient mutations are located at the interface of the BBS1 β-propeller and its helical insertion with the N-terminus of BBS4 (R160Q, E224K, R268P, L288R), underlining the importance of this interface. **(D)**: The mutation N309K disturbs a tight interaction of BBS4 with the main-chain of BBS18 in the “U-bolt”-region. L58^*^ eliminates the C-terminal helix of BBS18 but leaves the “U-bolt” region that clamps together BBS4 and BBS8 intact.

### BBS18 is a central stabilizing component of the core BBSome

From co-expression and pulldown studies of the individual BBSome subunits, we know that BBS18 is important for the stability of larger BBSome subcomplexes ^26^. The cryo-EM structure of the core BBSome delivers an explanation for these findings, revealing that BBS18 is involved in a large amount of stabilizing interactions with the TPR domains of BBS4 and BBS8 (Figure 3A). Because of its unfolded U-bolt region, BBS18 forms a large interaction surface with these subunits, resulting in high solvation free energies (Δ^i^G) (as analyzed by the Pisa server ^28^) (Figure 3A, Table S2).

The short helix of BBS18 (residues L61 to N80) protrudes from the BBSome core complex and only loosely interacts with the GAE domain of BBS1 (Figure 3B) and with a loop of BBS8 (Figure S2B). Despite this limited interaction with other subunits, the helix is important for the proper assembly of the BBSome as has been shown previously, analyzing a disease-causing null mutation in BBS18 (L58^*^) which results in the loss of the helix ^29^.

Taken together, these findings suggest that BBS18 functions as a structural scaffold protein that is essential for the proper assembly and structural stability of the BBSome complex.

### Surface complementary β-propeller-TPR interactions

Interestingly, both structurally analogous TPR containing subunits BBS4 and BBS8 interact with β-propellers of BBS1 and BBS9, respectively. In both cases, it is the N-terminus of the TPR containing subunits that binds to the central grooves of the β-propellers (Figure 3C,D), the mode of interaction, however, differs.

As previously described ^24^, the β-propeller of BBS9 is negatively charged around its central cavity. Our structure reveals that it interacts with a positively charged patch of BBS8 (Figure 3C). The surface charges at the center of the β-propeller of BBS1 and the N-terminus of BBS4, however, are more evenly distributed and the interface at this position is mostly stabilized by hydrophobic interactions (Figure 3D). In addition, the superhelically arranged TPRs of BBS4 and BBS8 wind around and extensively interact with the side of the respective β-propeller. These interfaces are mostly stabilized by potential ionic interactions (Figure 3C,D). Moreover, the interface between BBS8 and the side of the BBS9 β-propeller contains two pairs of methionines, which potentially interact via hydrophobic and S/π interactions (Figure 3C).

Mutations at the BBS4-BBS1 and BBS8-BBS9 interfaces (Table 1), such as Q325R ^25^ at the side of the BBS9 β-propeller (Figure 4B) or E224K ^30^ and R268P ^31^ in the central cavity of the BBS1 β-propeller (Figure 4C) lead to disease in patients, indicating that proper interaction between these subunits is crucial for a functional BBSome.

**Table 1:** Disease-causing variants at the interface between BBSome subunits. Only mutations that sit at the interface between BBSome subunits and likely have an influence on the stability of the complex have been analyzed.

| **BBS gene** | **mutation** | **phenotype** | **Reference** |
| --- | --- | --- | --- |
| BBS1 | L518P | BBS | {Mykytyn:2003fo} |
|  | N524Δ | BBS | {Deveault:2011eu} |
|  | R160Q | BBS, RP | {Sharon:2015vc, Deveault:2011eu} |
|  | E224K | BBS | {Redin:2012fh} |
|  | R268P | BBS | {EstradaCuzcano:2012dv} |
|  | L288R | BBS | {Muller:2010in} |
| BBS4 | N309K | BBS | {Muller:2010in} |
| BBS8 | Q439H | BBS, RP | {Ullah:2017vi, Goyal:2016be} |
|  | Q445K | RP | {vanHuet:2015wq} |
| BBS9 | Q325R | BBS | {Chou:2019gb} |
| BBS18 | L58* | BBS | {Scheidecker:2014dg} |

In contrast to BBS4, BBS8 contains a long loop between tyrosine 53 and lysine 180, connecting the first N-terminal TPR domain with the rest of the TPR superhelix. This loop is not fully resolved in our structure, but can be partly visualized at lower resolution (Figure S2A-B). Starting after the first TPR domain, it winds through the center of the complex and back to the following TPR domains. It interacts with BBS18 and with several TPRs of BBS8 and does not interrupt the TPR superhelix (Figure S2B). Interestingly, the sequence of the start and end region of this loop varies in different human BBS8 splice variants, expressed in different tissues ^32^. For example, Exon 2a of BBS8 is only expressed in retina, resulting in an additional insert of 10 residues at the beginning of the loop. Mutations inducing the splicing of Exon 2a result in the expression of BBS8 without these 10 additional residues. Although the sequence of the resulting loop is similar to the one of the loop in other tissues, it results in non-syndromic retinitis pigmentosa ^32^, suggesting that this loop is involved in a tissue-specific essential function.

### The C-terminus of BBS9 forms a GAE binding motif

A prominent feature in the core BBSome structure is the expanded domain arrangement of BBS1 and BBS9 which allows the wrapping of these subunits around the central BBS4 and BBS8 subunits (Figure 2C,D). BBS1 and BBS9 interact with each other via their C-terminal domains (GAE, platform, and α-helical domain) in such a way that the β-propellers orient to the opposite direction locating to the periphery of the complex where they interact with the TPR domains of BBS4 and BBS8 (Figure 2C,D, Figure 4A). The interaction of BBS1 with BBS9 is essential for the function of the BBSsome, as mutations L518P ^33^ and N524Δ ^34^ at the interface between the GAE domain of BBS1 and BBS9 are pathogenic (Figure 4A, Table 1). The GAE domain of BBS1 is also involved in interactions with BBS8. The glutamine residues 439 and 445 in BBS8 are located at this interface and their mutation results in disease (Q439H, Q445K) ^25,35–37^(Figure 4A, Table 1).

The GAE domains of BBS1 and BBS9 interact strongly with each other (Figure 4A). Since their structure is very similar, it can be imagined that these domains would induce self-dimerization of either BBS1 or BBS9. Indeed, in previous studies ^24, 26^, it was shown that the isolated C-terminal region of BBS9, including the GAE domain, forms a dimer in solution. A superposition of BBS9 monomers reveals that an interaction via their GAE domains would not result in steric clashes (Figure S2C). Taken together with our observation that the GAE strongly interact in the BBSome core complex, we propose that the strong heterodimeric interaction between the GAE domains is the reason for isolated BBS9 to homodimerize via its GAE domain.

### BBSome subcomplexes bind to phosphoinositides in the absence of BBS5

Compared to the other subunits, BBS5 is more loosely attached to the BBSome core complex as we could only identify the subunit in a subpopulation of particles (Figure S1, S4). In addition, the resolution is limited in this region, indicating a high degree of conformational flexibility.

BBS5 is composed of two pleckstrin homology (PH) domains that were predicted to be close structural relatives to the PH-like domains PH-GRAM ^38^ and GLUE ^39^. In the BBSome core structure, the PH domains are laterally rotated by 90° to each other (Figure 2E). Like PH-GRAM and GLUE, BBS5 was shown to bind phosphoinositides, with the highest preference for phosphatidylinositol 3-phosphate (PI(3)P) and phosphatidic acid (PA), which was suggested to be crucial for ciliogenesis ^40^. Besides BBS1 that indirectly interacts with membranes via Arl6 ^16, 23^, BBS5 likely mediates the contact to membranes by direct interactions with phosphoinositides ^40^. The specific binding to certain PIPs is a potential way to regulate BBSome transportation in and out of the cilium, as the composition of ciliary membranes differs from that of the plasma membrane ^41–43^.

To find out whether BBS5 is the only subunit of the BBSome complex that interacts with phosphoinositides, we compared the binding of different phosphoinositides to the core BBSome complex containing BBS 1, 4, 5, 8, 9 and 18 with a smaller complex that lacks BBS5 and BBS1 (Figure 6A-C). We found that the core BBSome complex interacts preferably with PI(3)P, PI(3,5)P_2_, PI(4,5)P_2_, PI(5)P, and PA. This pattern is similar to the one observed for BBS5 alone 40, however, the interaction with PI(3,5)P2 and PI(4,5)P_2_ is more pronounced in the case of the BBScome core complex (Figure 6B). Surprisingly, the smaller 4mer complex also binds specifically to the same phosphoinositides as the BBS5 and BBS1-containing complex and especially strong to PA (Figure 6A). This shows that BBS5 is not exclusively responsible for phosphoinositide and PA binding and that respective binding sites must exist on one or more of the other subunits, namely BBS 4, 8, 9 or 18.

### Membrane association and cargo recognition of the core BBSome

The BBSome is recruited to membranes by the small GTPase Arl6 in a GTP-dependent manner ^16^. A crystal structure of Arl6 with the β-propeller of BBS1 from *Chlamydomonas reinhardtii* revealed how Arl6 interacts with the BBSome in atomic detail ^23^. To analyze how the core BBSome would orient on the membrane, we placed Arl6 at the same position as in the crystal structure by overlaying the respective BBS1 β-propeller domains (Figure 6D). The BBS1 β-propeller is located at the periphery of the core BBSome complex, and – in the absence of BBS2 and BBS7 - is freely accessible to bind to membrane-attached Arl6. Such an orientation would position a positively charged surface patch of the core BBSome close to the membrane, which is favorable for interactions with the negatively charged membrane surface. Importantly, this orientation also leaves a negatively charged cleft in the BBSome structure oriented perpendicular to the plane of the membrane (Figure 6E,F). We have previously found that prominent features of GPCR cargo molecules which determine binding to the core BBSome are motifs composed of aromatic and basic residues, many of which are found in the third intracellular loop and the C-terminal tail of GPCRs ^26^. The negatively charged cleft within the BBSome probably interacts specifically with these positively charged motifs and is thereby directly involved in cargo recognition (Figure 6E-G). This would also position the motifs close to BBS1, a subunit which was shown to be particularly important for cargo recognition ^16–22^.

The negative charge in the cleft is not formed by BBS1 alone, but also by residues of all other core BBSome subunits, except BBS5. There are three “hotspots” of negative charge. One of them is located at the contact surface of the GAE domain of BBS1 and the 5α domain of BBS9 (Figure 5B,C), the second one is deeply buried in the core BBSome complex at the junction point where the BBS18 U-bolt ends and descends into a short α-helix (Figure 5D,E), and the third one is determined by the α-helical insertion within the BBS1 β-propeller, which contains many glutamate or aspartate residues (Figure 5F,G). The α-helical insertion can only get in contact with cargo peptides that extend far into the negatively charged cleft (Figure 6D). The β-propeller itself and the C-terminal GAE domain of BBS1 are also accessible from within the cleft. They have a more balanced charge distribution and might contribute to cargo binding by hydrophobic and ionic interactions at the base of the cleft, which would explain the high importance of BBS1 for cargo interactions.

**Figure 5:**
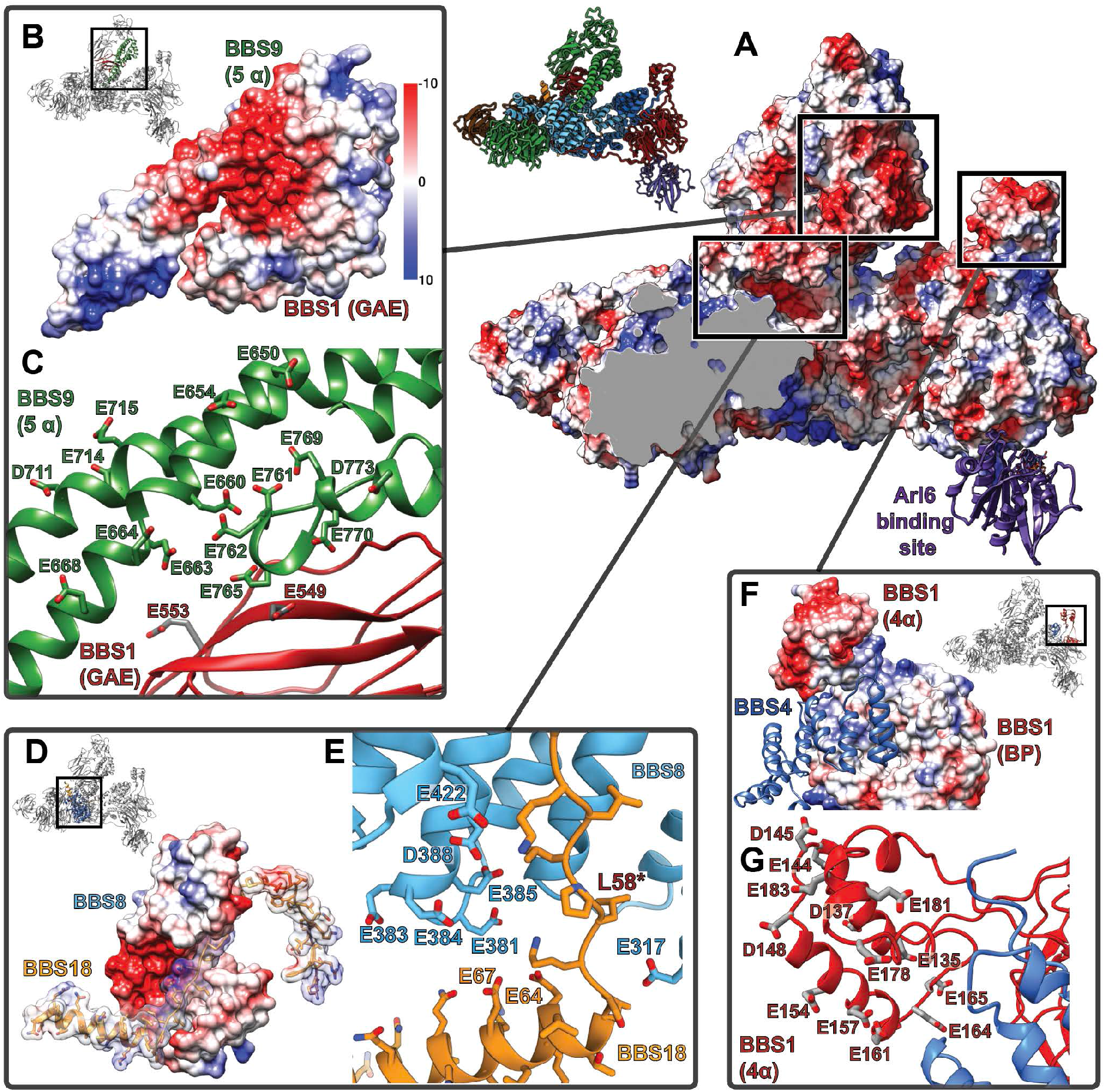
Highly negatively charged regions in the central cleft of the BBSome. **(A)**: An open cleft within the center of the core BBSome contains multiple highly negatively charged regions that might be involved in cargo binding. The Arl6 binding site, as deduced from the crystal structure of the β-propeller of BBS1/Arl6 23, is shown as purple ribbon. **(B,D,F)**: Surface charges in three hotspots of negative charge within the central cleft. **(C,E,G)**: Positions of negatively charged residues.

**Figure 6:**
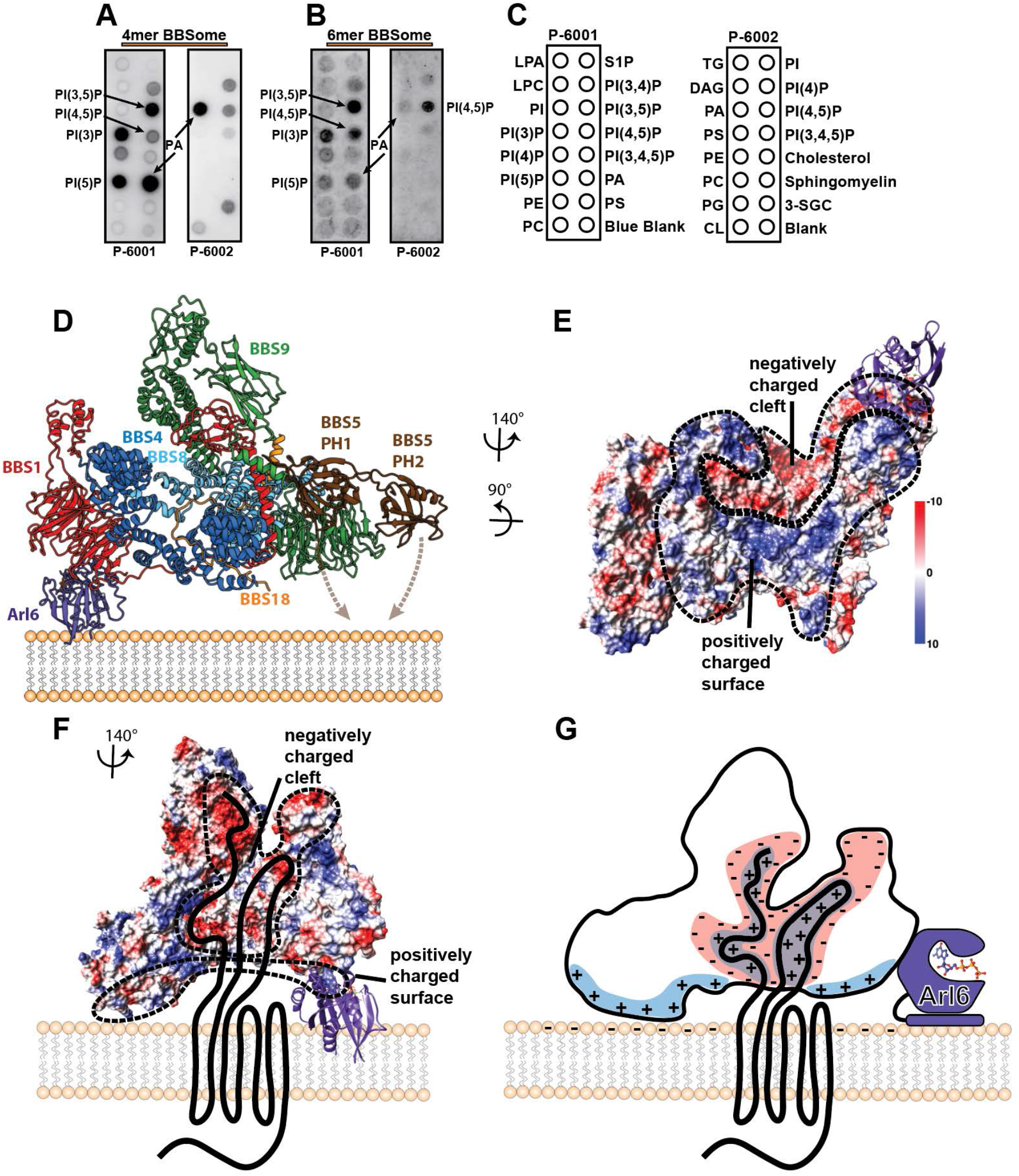
Interaction of the core BBSome with membranes. **(A-C)**: PIP strip experiments indicate that BBSome subcomplexes interact specifically with PIPs even in the absence of both the Arl6-binding subunit BBS1 and the previously described PIP-binding subunit BBS5 40. **(D)**: Potential orientation of the BBSome core complex towards the membrane. The orientation of Arl6 towards the BBSome was deduced from the crystal structure of the β- propeller of BBS1/Arl6 23, which was overlaid with the BBS1 β-propeller from the BBSome core complex. In such an arrangement, a positively charged surface of the core complex would be oriented towards the membrane **(E,G)**and a negatively charged cleft in the vicinity of BBS1 is favorably positioned to accept BBSome-binding regions from cargo proteins like GPCRs **(E-G)**, which were found to be mostly positively charged 26. A model how GPCRs might be recognized by the BBSome is depicted in **(G)**.

The large size of the binding cleft with different “hotspots” suggests that cargo recognition is likely very complex and variable, involving the interaction with different BBSome subunits. The proper study of BBSome-cargo interactions therefore requires the full BBSome complex as cargo binding to single subunits, albeit relevant, does not take interaction to multiple sites into account. For example, we previously identified a peptide fragment from the C-terminal part of SSTR3 which binds to the core BBSome with ~100-fold higher affinity than to the isolated BBS1 β-propeller (aa1-430) ^26^.

Since the BBS5 and Arl6 binding sites are located on opposite sides of the core BBSome complex (Figure 6D), a simultaneous membrane binding of both motifs would require a curved membrane surface, as it is present on the inner cilium wall, and/or conformational rearrangements of the complex. The observed dynamic properties of BBS5 (Figure S1, S4) are probably the prerequisite for the subunit to be able to reorient and position one or both of its PH domains closer to the membrane, thereby enabling interaction with the membrane.

While this manuscript was being prepared, the structure of the bovine BBSome complex at 4.9 Å was published, in which the Arl6 binding site is blocked by the subunits BBS2 and BBS7 ^25^. These two additional subunits form a highly intertwined lobe that contacts the BBSome core at the C-terminal hairpin in BBS9 and at the β-propeller of BBS1, thereby temporarily blocking the binding site for Arl6 ^25^. The overall architecture of the two complexes is very similar, but interestingly, in a rigid body overlay of our structure with the bovine BBSome, the Arl6 binding site does not clash with BBS2 or BBS7 (Figure S6). This is due to a different orientation of the BBS1 β-propeller to which Arl6 binds. In the structure of the bovine BBSome complex the BBS7 β-propeller interacts with the β-propeller of BBS1 and thereby pulls it towards the BBS2-BBS7 lobe. The small change of the BBS1 β-propeller orientation results in a ~20° change in the angle of Arl6 binding to the complex (Figure S6F-H). We therefore believe that our BBSome core structure represents the open conformation of the complex in contrast to the autoinhibited closed state of the bovine BBSome complex.

The additional subunits BBS2 and BBS7 extend the interaction surface that we suggest to bind to ciliary membranes (Figure 6E, Figure S6C), and narrow down the opening to the negatively charged cleft. However, it is still sufficiently large for peptides from cargo proteins to enter the cleft (Figure 6G,H, Figure S6C).

It is interesting to note that in native and heterologous expression, the top lobe formed by BBS2 and BBS7 readily dissociates ^25^ from the complex or renders it insoluble ^26^. Likewise, BBS5 was present in substoichiometric amounts in both purifications. Chou *et al.* suggested that the stability of the α-hairpin in the C-terminal domain of BBS9 likely depends on the stabilization by its partner hairpin in BBS2. We found that in the absence of BBS2, the BBS9 hairpin folds towards BBS8 and is stabilized that way (Figure S6D,E). This further supports our hypothesis that the core BBSome is an independently stable entity ^26^, and that BBS2 and BBS7 is probably not required in every step within the lifecycle of the BBSome.

## Conclusion

Future studies should reveal the details of how cargo proteins get recognized and how the “top lobe” formed by BBS2 and BBS7 makes space for cargo binding, particularly in the context of BBSome interactions with Arl6 and the IFT complex. For this it will be crucial to obtain structures of the BBSome complex bound to different cargo peptides or full cargo proteins. The architecture of the extended negatively charged cleft within our structure of the core BBSome suggests that cargo binding depends on the 3-dimensional arrangement of all BBSome subunits, and that different cargo proteins might utilize different binding modes within the cleft. This would also suggest that the BBSome complex binds differently to cargo proteins in its open or its closed conformation.

Other intriguing questions to be addressed in the future include determining the precise orientation of the BBSome on membranes, and the relevance of interactions with phosphoinositides via BBS5 or via currently unidentified phosphoinositide binding motifs on one of the subunits BBS4, 8, 9 or 18 (compare Figure 6A-C).

## Materials and Methods

### Purification of BBSome subcomplexes

The core BBSome (BBS 1, 4, 5, 8, 9, 18) and the 4mer BBSome subcomplex (BBS 4, 8, 9, 18) were cloned and purified as described previously ^26^. Briefly, the proteins were overexpressed in Hi5 insect cells (Thermofisher Scientific), harvested by centrifugation at 3000 rpm, and resuspended in lysis buffer (50 mM Hepes pH 8.0, 150 mM NaCl, 5 mM MgCl_2_, 10% glycerol, 1 mM PMSF, 1 mM benzamidine, 1 mM TCEP). The resuspended cells were lysed with a Dounce Tissue Grinder (Wheaton, Millville, NJ), cell debris was removed by centrifugation at 25.000 rpm, and the supernatant was used for further protein purification.

For affinity purification, the cell supernatant was loaded on a column filled with Strep-Tactin Superflow high capacity resin (IBA). The column was washed and the proteins were eluted with elution buffer (50 mM Hepes pH 8.0, 150 mM NaCl, 5 mM MgCl_2_, and 10% glycerol, 1 mM TCEP, and 10 mM D-desthiobiotin (IBA)). The eluted protein complexes were further purified by Ni^2+^–NTA (Qiagen, Germany) affinity and/or anti-Flag M2 affinity chromatography (Sigma, Germany). Affinity-purified proteins were subjected to size-exclusion chromatography on a Superdex 200 10/300 column or on a Superose 6 5/150 column (GE Healthcare, Germany) in gel filtration buffer (50 mM Hepes pH 8.0, 150 mM NaCl, 5 mM MgCl_2_, 10% glycerol and 0.1 mM TCEP).

### Protein-Lipid Overlay assays

The affinity of the core BBSome complex (BBS1, 4, 5, 8, 9, 18) and the 4mer BBSome complex containing BBS4, 8, 9 and 18 to different lipids immobilized on hydrophobic membranes (“PIP-strips” P-6001, P6002, Echelon) were probed in a protein-lipid overlay assay according to the manufacturer’s instructions. In brief, PIP-strips were blocked with TBS-T + 3% fatty acid–free BSA and then incubated with 7.5 μg/ml complex for one hour at room temperature. After washing three times with TBS-T + 3% fatty acid–free BSA, immobilized complexes were detected by a western blot against the Flag-tag on BBS8.

### Chemical crosslinking to improve complex stability

To improve the stability of the core BBSome complex for cryo-EM experiments, the purified protein was diluted to 0.1 mg/ml in gel filtration buffer and treated with 0.05% glutaraldehyde at 20 °C for two minutes. The reaction was stopped by adding TRIS buffer to a concentration of 100 mM. The protein was then concentrated and subjected to size-exclusion chromatography on a Superose 6 5/150 column (GE Healthcare, Germany) in plunging buffer (20 mM TRIS-HCl pH 7.5, 150 mM NaCl and 0.1 mM TCEP) to remove particles that aggregated during the crosslinking procedure. Fractions containing the crosslinked BBSome complex were immediately used for cryo-EM experiments.

### Sample vitrification

The concentration of the core BBSome as derived from chemical crosslinking was adjusted to 0.05-0.08 mg/ml and 4 μl of sample was immediately applied to a glow-discharged UltrAuFoil® R1.2/1.3 holey gold grid (Quantifoil). After two minutes incubation at 13 °C and 100% relative humidity (RH), the sample was manually blotted, replaced by fresh 4 μl sample, and then automatically blotted and plunged in liquid ethane using a Vitrobot (FEI).

### Electron microscopy and image processing

Cryo-EM datasets were collected on a Titan Krios electron microscope (FEI) equipped with a post-column energy filter, a Volta phase plate (VPP) and a field emission gun (FEG) operated at 300 kV acceleration voltage. A total of 15,266 micrographs were recorded on a K2 direct electron detector (Gatan) with a calibrated pixel size of 1.07 Å. The in-column energy filter was used for zero-loss filtration with an energy width of 20 eV. In total 50 frames (each 300 ms) were recorded, resulting in a total exposure time of 15 s and a total electron dose of 67 e^−^Å^−2^. Data was collected using the automated data collection software EPU (FEI), with a defocus range of −0.3 to −1.0 μm. The position of the VPP was changed every 60 to 120 images, resulting in phase shifts of 30-120 degrees in > 95% of all micrographs. Beam-induced motion was corrected for by using Motioncor2 ^44^ to align and sum the 50 frames in each micrograph movie and to calculate dose-weighted and unweighted full-dose images. CTF parameters were estimated from the unweighted summed images and from micrograph movies utilizing the “movie mode” option of CTFFIND4 ^45^. For subsequent steps of data processing using the software package SPHIRE/EMAN2 ^27^, dose-weighted full dose images were used to extract dose-weighted and drift-corrected particles with a final window size of 280 × 280 pixels.

A total of 10 datasets were collected and were successively processed by a combination of manual and automated particle extraction using crYOLO ^46^, 2D sorting using the iterative stable alignment and clustering (ISAC) as implemented in SPHIRE, and merging with existing data to consecutively improve the size and quality of the derived particle stack. An initial model for 3D refinement was generated from the ISAC 2D class averages using RVIPER from SPHIRE, and was used as input for the first 3D refinement using MERIDIEN (3D refinement in SPHIRE). The obtained 3D reconstruction of MERIDIEN was then sharpened and filtered to its nominal resolution, and used as input for subsequent 3D refinements. Completeness of data collections was evaluated based on the improvements in resolution of the resulting final reconstructions, which plateaued at 4 Å resolution from a cleaned set of 724,828 particles, as estimated by the ‘gold standard’ criterion of FSC = 0.143 between two independently refined half maps.

To further improve the quality of the derived particle stack, an alternative processing attempt was tested in which all 15266 micrographs were combined and subjected to automatic particle picking using crYOLO, selecting 1,973,261 particles with a high confidence score (threshold 0.7) and additional 858,068 particles that had a lower score (threshold greater than 0.2 but smaller than 0.7). Both sets of particles were subjected to 2D sorting, and the best particles of each subset were merged and sorted again to obtain a new particle stack of 884,341 particles. After 3D refinement, the final reconstruction had a slightly improved quality compared to the first processing attempt. However, it was also apparent that a significant number of “good” particles were only selected in one of the two sorted particle stacks, i.e. indicating that about 30% of high-quality particles were lost during refinement. By combining the two particle stacks from both independent processing attempts and removing all duplicate particles, we rescued about 220,000 “good” particles that were sorted out in one or the other processing attempt. The derived combined stack of 1,103,959 particles could thus be subjected to a final, more stringent 2D sorting round to obtain a final particle stack with 862,114 particles that, after 3D refinement, resulted in a 3.8 Å reconstruction with significantly improved quality compared to the individual refinement attempts.

3D clustering using SORT3D of the SPHIRE suite was performed with a 3D-focussed mask that includes BBS5, which was first apparent in a variability map of the full particle stack as calculated using 3DVARIABILITY from SPHIRE (Figure S4). 3D clustering separated the particles into four 3D clusters which were then subjected to local 3D refinement using MERIDIEN. One of the generated 3D reconstructions clearly indicated BBS5 density, resulting in a 4.3 Å resolution map of the core BBSome with BBS5 (Figure S4). An analysis of the 3D variability in this map indicated no remaining variability in the BBS5 focus region, while the variability in other regions of the complex remained similar as in the full particle stack (e.g. indicating some conformational heterogeneity around the 4α helical insert in the β-propeller of BBS1, around the platform domain of BBS9, at the tip of the hairpin within the 5α helical domain of BBS9, and at the unresolved C-terminal end of BBS4).

### Model building

For building an atomic model of the BBSome core comples, we used the two available crystal structures of the β-propeller domains of *C. reinhardtii* BBS1 ^23^ and of human BBS9 ^24^ as starting points for the assignments of BBS domains. For this, the human homology model to the *C. reinhardtii* BBS1 β-propeller was generated using HHpred ^47^, and the structure of a helical insert in the β-propeller (Pro127-Gln197) that is missing in the crystal structure was predicted using RaptorX ^48^. For the human BBS9 β-propeller, the pdb entry 4YD8 could be directly used as starting model after minor modifications (i.e. changing the selenomethionine residues to methionine and adding a few loops that were missing in the crystal structure). The large helical insert in BBS1 that is not present in BBS9 allowed an unambiguous placement of both structures into the density. BBS4 and BBS8 are the two subunits that are predicted to fold into TPR repeats, and from biochemical data it is apparent that BBS9 forms a stable binary subcomplex with BBS8 ^26^. Since in the cryo-EM map two superhelical arrangements of TPR motifs are visible and BBS9 forms no interactions with one of them, the correct placement of BBS9 and BBS1 also allowed an unambiguous assignment of BBS4 and BBS8. Initial models for BBS4 and BBS8 were generated by *de novo* structure prediction using RaptorX ^48–51^ followed by flexible fitting of fractions of the RaptorX structure predictions into consistent parts of the density using iMODFIT ^52^ and subsequent manual model correction in Coot ^53^. The validity of the fit could be verified based on side chain consistency with the map as BBS4 and BBS8 are positioned in a central region of the complex with sufficiently high local map resolution. Consistent with the RaptorX prediction, we observed that a region of about 130 residues in BBS8 (Tyr53-Lys180) is only partially structured and forms an extended loop that winds through the center of the complex and back (Figure S2A,B).

The assignments of the C-terminal parts of BBS1 and BBS9 were less obvious as they are separated by partially flexible linkers from the N-terminal β-propeller domains. We could assign the GAE domains of BBS1 and BBS9 based on the clearly resolved connection of the BBS9 GAE domain to the C-terminal platform and α-helical domains that are missing in BBS1. Furthermore, the connection of the BBS1 GAE domain could be traced up to a small gap of about 20 residues to its corresponding N-terminal β-propeller, which would not allow an alternative assignment. The structures of all these domains were also predicted with RaptorX, flexibly fitted with IMODFIT and manually corrected with Coot as described above. The resolution of the GAE domains was sufficient to verify the validity of the model based on side-chain densities, and even in the lower resolved part of the platform and α helical domains of BBS9, multiple hallmark residue side chains allowed a clear validation of the overall fold, although some solvent-exposed loops are only weakly defined.

After placing BBS1, 4, 8 and 9, a prominent stretch of well resolved density remained unexplained that winds through the superhelices formed by BBS4 and BBS8 and connects to a short α-helix. With a total length of about 60 residues and no obvious covalent connection to one of the other subunits, we were confident that this stretch represents a major part of the 93 residue subunits BBS18, which is also consistent with biophysical data showing that BBS18 forms stable subcomplexes with BBS4 and with BBS4, 8 and 9 ^26^. With few initial indications how to assign the primary sequence to this linear, mostly unfolded domain, we utilized the following method to find the correct frame: We first built a 59 residue poly-alanine model, then assigned all 33 possible frames of BBS18 sequence to this model, and further refined each of them automatically using Phenix real-space refinement ^54^(Figure S5A-E). Comparing the correlation of the map to the refined models, we observed a clearly separated best hit for one sequence assignment (Figure S5A,E) with no obvious discrepancies between map and model, while the other models showed significant deviations from map to model that could not be explained by noise, imperfect modelling or a lack of resolution. The final model was further manually refined in Coot and corresponds to residues Val26-Gln80 of BBS18 (Figure S5F).

After placing all subunits except BBS5, missing residues that were visible in the density were placed and all subunits were further manually refined in Coot. Atom clashes were removed by energy minimization using PHENIX ^54^, followed by another round of manual refinement. Compared to the other subunits of the core BBSome, BBS5 is more loosely attached at the periphery of the complex, and could only be identified in a subpopulation of particles that was isolated by 3D clustering (Figure S1,S5). The best 3D class containing BBS5 allowed a reconstruction with an average resolution of about 4.3 Å (Figure S1D, S4). The two PH domains of BBS5 were modelled with RaptorX and were positioned into the derived BBS5 density using Chimera (UCSF) *55*, followed by relaxation into the density in Coot (Figure S1E). An overlay of the maps derived from the full particle stack with the 3D cluster containing BBS5 allowed to position BBS5 relative to the higher resolved reconstruction from all particles in which BBS5 was not visible. BBS5 was then combined with the higher resolved full particle reconstruction to generate a combined molecular model of the human BBSome core complex.

### Visualization

Visualization, analysis and figure preparation was done with Chimera (UCSF) ^55^. Local resolution gradients within a map were estimated using LOCRES as implemented in the SPHIRE suite, and final densities were filtered according to the calculated local resolution unless stated otherwise. Resolution gradients were visualized by coloring the corresponding maps according to the local resolution in Chimera (Figure S3D). To visualize surface electrostatic potentials, the correct protonation state of the core BBSome was predicted using the H++ web server (http://biophysics.cs.vt.edu/H++^56^) and the hydrogenated model was colored according to its electrostatic potential in Chimera. 3D average and variability maps were calculated using 3DVARIABILITY of the SPHIRE package and visualized in Chimera. Angular distribution plots were generated using PIPE from SPHIRE. In addition to the binned 2-D class averages produced by ISAC that were used for the particle selection process, unbinned class averages showing high-resolution features were calculated using COMPUTE_ISAC_AVG from SPHIRE for visualization purposes.

## Acknowledgements

We thank Dr. Daniel Prumbaum for his support during EM data collection, the SPHIRE developer team for support in image processing and the whole Raunser lab for support and fruitful discussions. This work was funded by the Max Planck Society (to SR).

## Author Contributions

S.R. and A.W. designed the project. B.U.K cloned, expressed and purified the BBSome complexes, performed the protein-lipid overlay assays, and prepared cryo-EM specimens. B.U.K and O.H. collected the EM micrographs. B.U.K performed the image processing, built the atomic models and prepared the figures. C.G. provided support for the cryo-EM workflow. B.U.K. and S.R. wrote the manuscript. All authors reviewed the results and commented on the manuscript.

## Supplementary Figures

**Figure S1:**
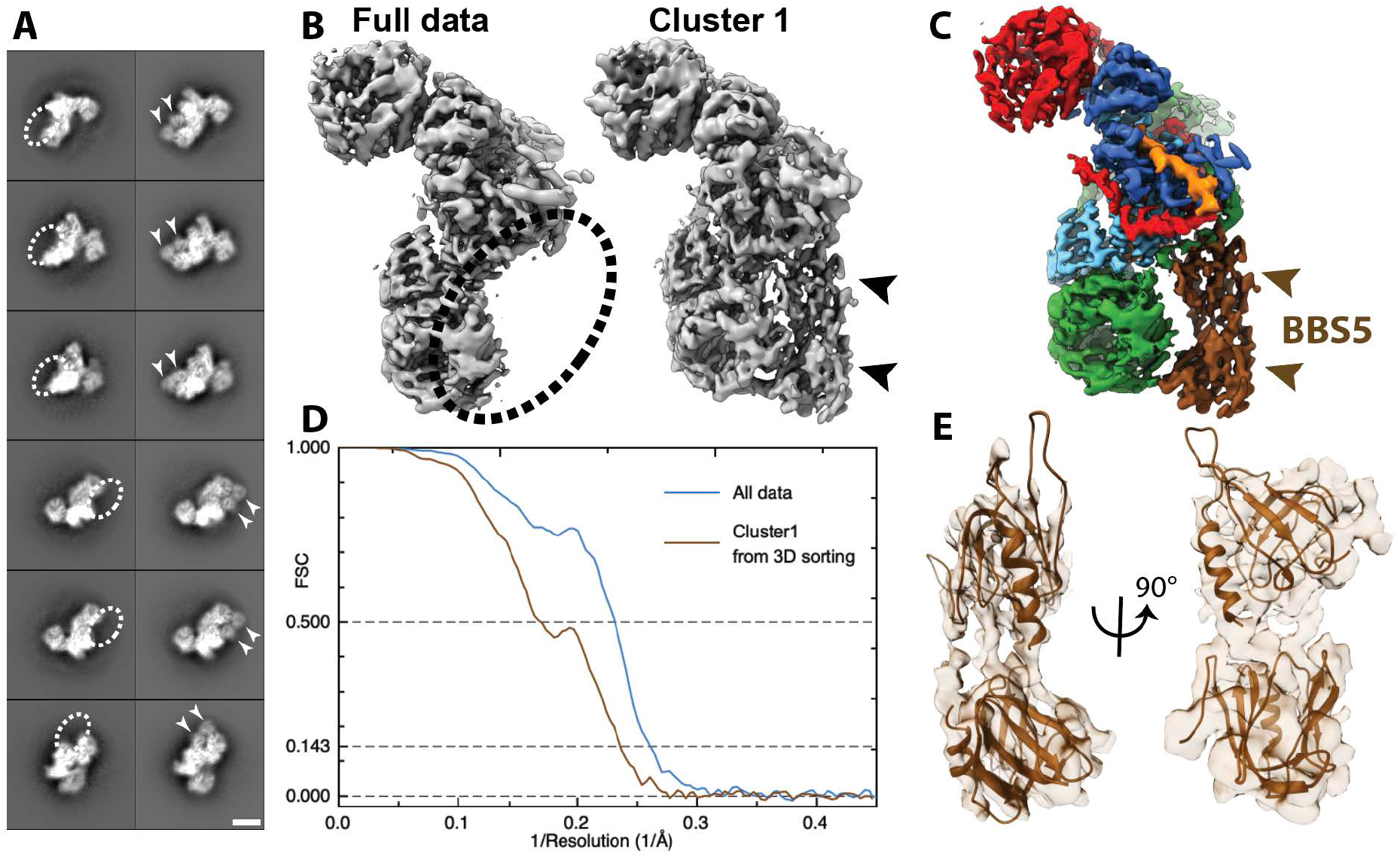
Positioning of BBS5 in the BBSome core complex. **(A)**: Pairs of exemplary 2D class averages of BBSome core complexes with and without BBS5 in different views (scale bar 5 nm) **(B)**: Comparison of the reconstructions from all available data and the best obtained cluster after 3D sorting which contains BBS5. **(C)**: The same view of the final, combined density map; **(D)**: FSC curves of the full dataset compared to the best cluster containing BBS5; **(E)**: fit of PH domains of BBS5 into density from Cluster1.

**Figure S2:**
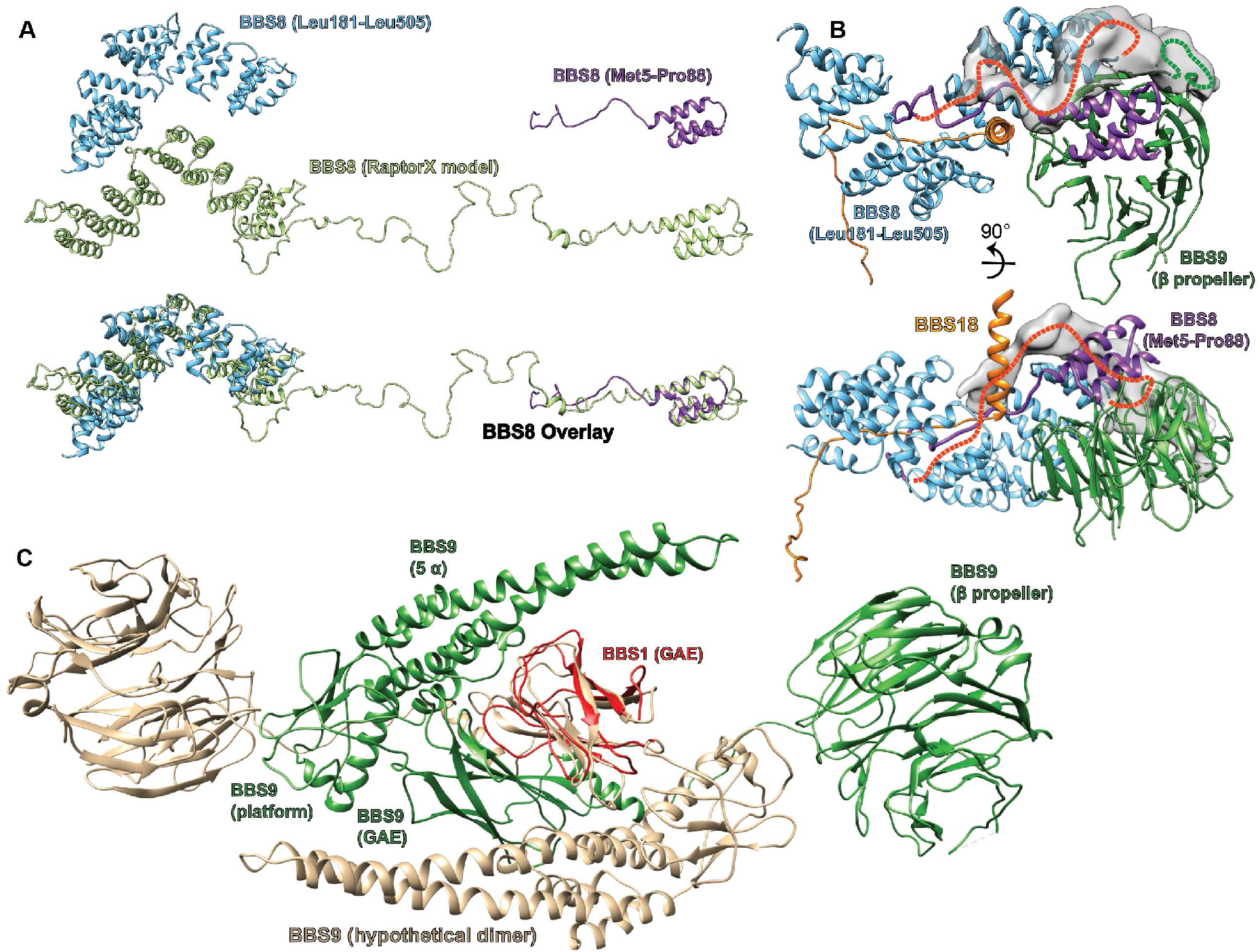
Partially structured insert in BBS8, and hypothetical model for BBS9 dimerization. **(A)**: A *de novo* structure prediction of human BBS8 as generated by Raptor X 48-51 indicates an unstructured region between a short N-terminal and a larger C-terminal TPR domain. This prediction is consistent with the final atomic model of BBS8 within the core BBSome structure, which contains both the N-and C-terminal TPR parts, but could only partially resolve the unstructured insert. **(B)**: The N-terminus of BBS8 (purple ribbon) descends into an unstructured region that could partially be traced to loop through the center of the core BBSome complex, forming interactions with BBS8 and the U-bolt region of BBS18. Extra density indicates how the loop approximately winds back to connect to the C-terminal part of BBS8, which is also indicated by a red dashed line. The loop seems to interact with an unresolved insert within the BBS9 β-propeller that also contributes to the unexplained density (green dashed line). **(C)**: Model for the dimerization of isolated BBS9 via its C-terminal GAE, platform and α- helical domains. A copy of BBS9 was superimposed with BBS1 via their homologous GAE domains, which provides a hypothetical dimerization interface of BBS9 that could explain why isolated BBS9 dimerizes via its C-terminus, while in the BBSome complex it does not dimerize 26.

**Figure S3:**
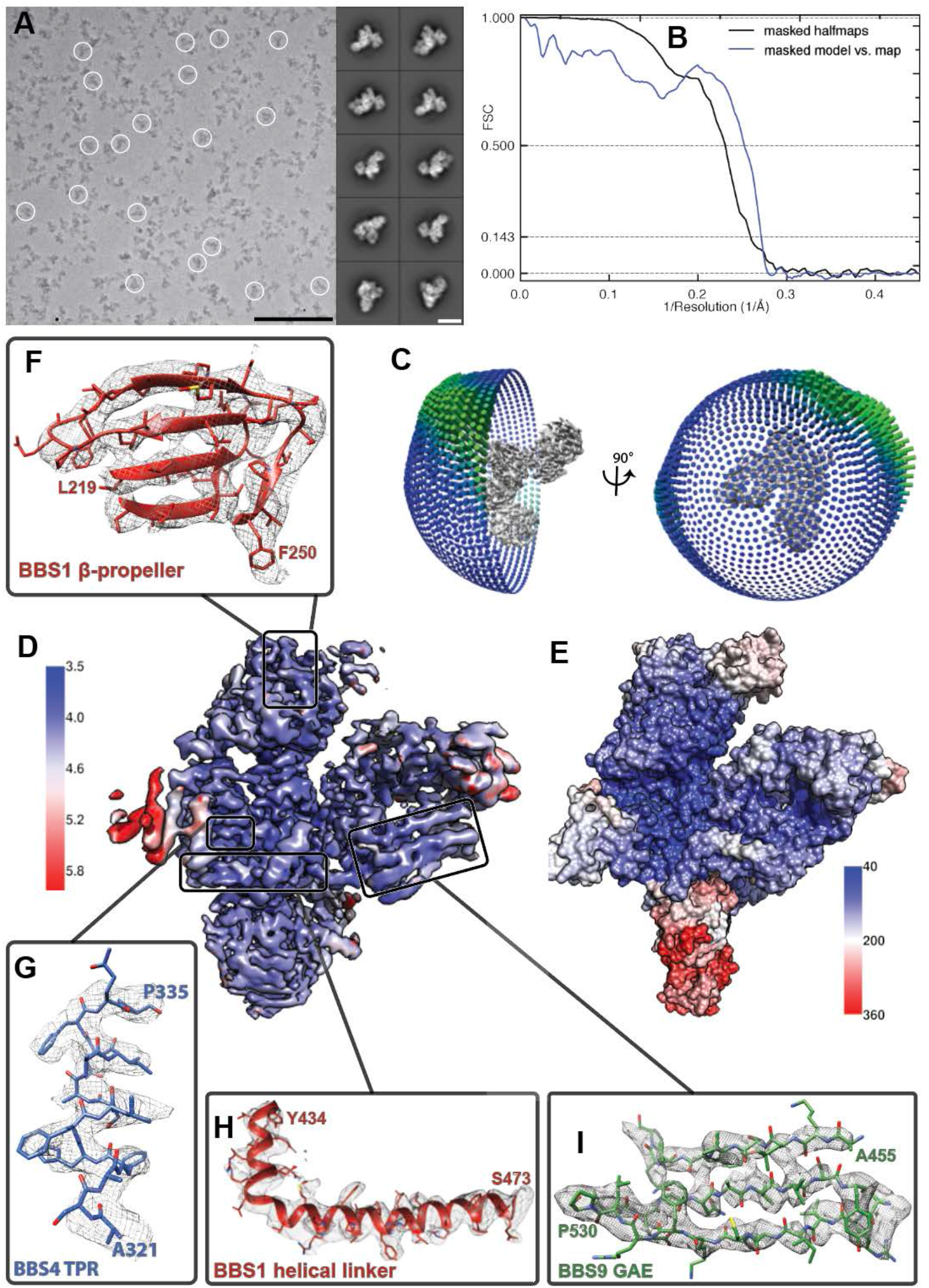
Quality of the model of the BBSome core complex. **(A)**: Cryo-EM micrograph and representative class averages of the core BBSome. Scale bars, 100 nm (micrograph) and 10 nm (class averages). **(B)**: FSC of two independently refined half data sets (black line), as well as the FSC between the volume calculated from all particles and the molecular model of the core BBSome (blue line). **(C)**: Angular distribution of particles that were used for the final reconstruction. **(D)**: Cryo-EM density map of the full dataset colored according to local resolution. **(E)**: Distribution of temperature factor (B-Factor) in the final model, including BBS5. The relative position and B-Factor distribution of BBS5 was derived from a reconstruction of a subset of particles (see Figure S1). **(F-I)**: Examples of the molecular model and the corresponding electron density map from different regions of the structure.

**Figure S4:**
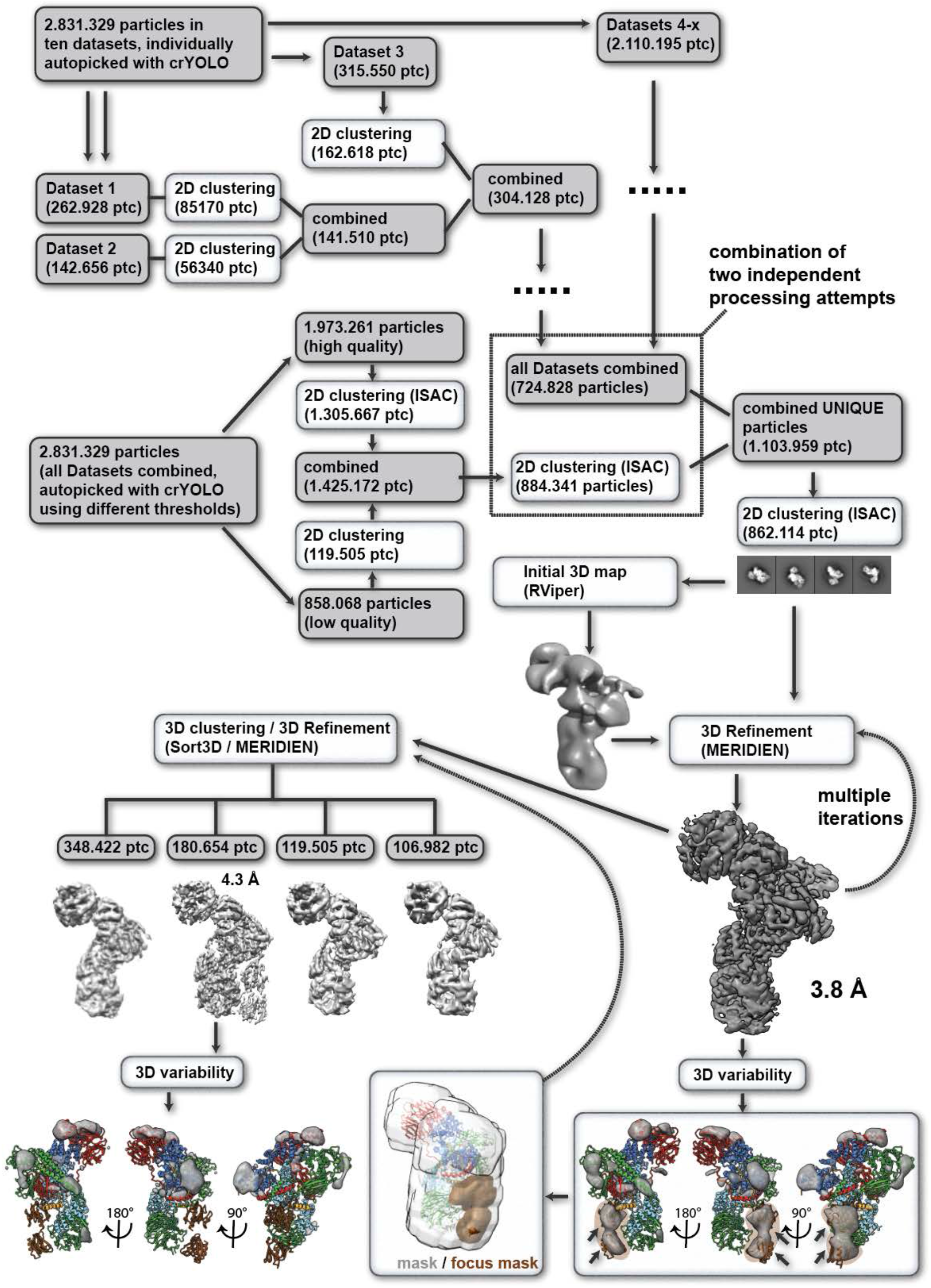
Single particle processing workflow for structure determination of the human core BBSome.

**Figure S5:**
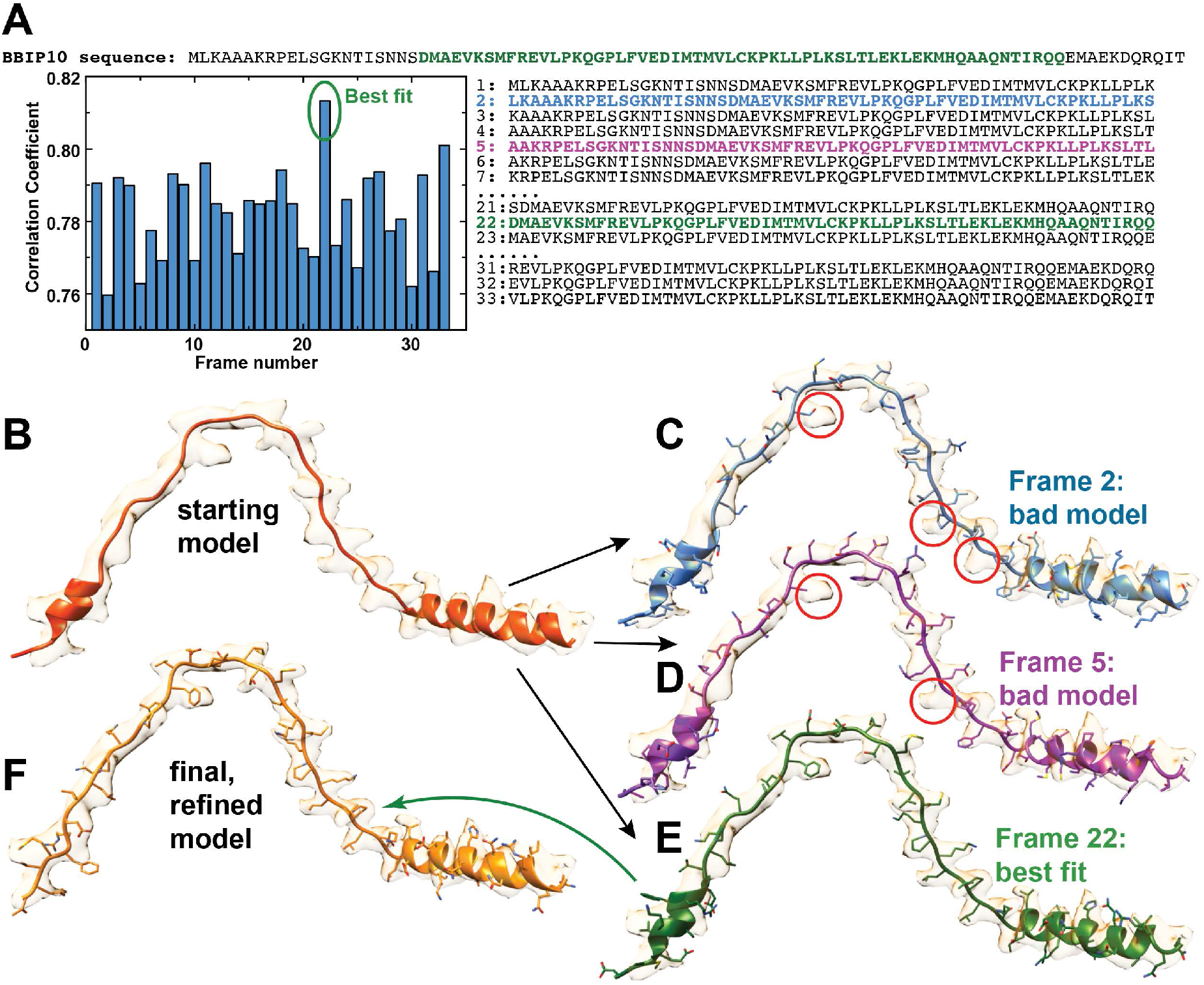
sequence assignment of BBS18. **(A)**: Only about two third of the 92 residues of BBS18 are visible in the calculated electron density, and besides a short helical region the protomer contains no defined secondary structure, which complicates an assignment of the correct sequence frame. To unambiguously find the correct frame, we first built a poly-alanine model into the density **(B)**. With only 92 residues of BBS18, there were 33 potential BBS18 sequence frames that could be assigned to the model. We generated all these 33 models, and further refined each of them using Phenix real-space refinement 54 **(C-E)**. The correlation of the map to the refined model showed a clear best fit for frame 22 **(E)**. Indeed, model #22 was in full agreement to the observed density. In contrast, the other models showed significant deviations from map to model that cannot be explained by noise, imperfect modelling or a lack of resolution (e.g. obvious side chain density for a residue that should not produce such density; **C,D**), We used model 22 for further manual refinement to obtain the final BBS18 model **(F)**.

**Figure S6:**
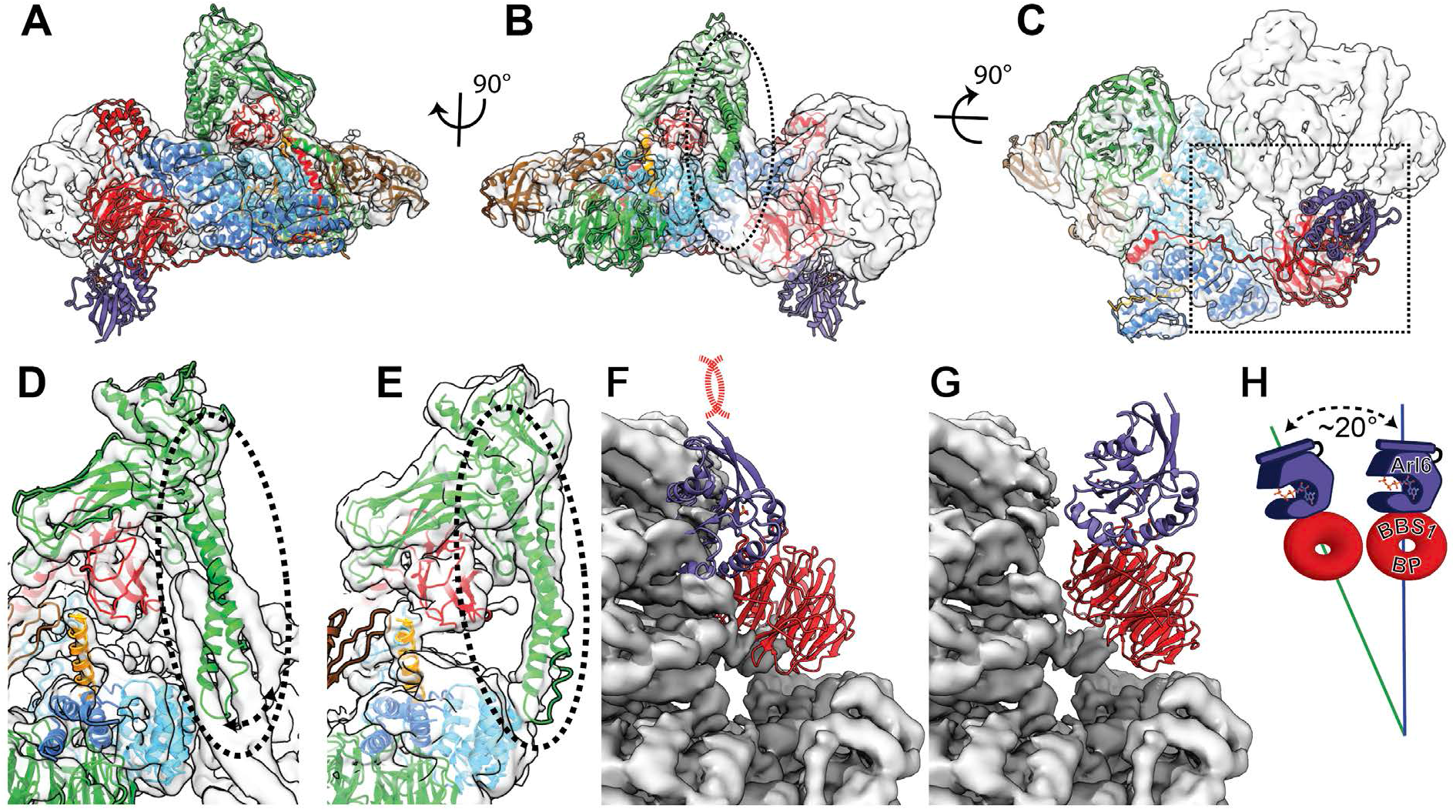
Comparison of bovine BBSome with human core BBSome. **(A-C)**: A rigid body fit of the human core BBSome structure into the bovine BBSome density 25 shows a similar overall arrangement, but distinct differences in the orientation of the α-hairpin of BBS9 in the C-terminal domain (dashed circle), and no clashes of the Arl6 binding site (purple domain). **(D-E)**: Closeup views of the α-hairpin of BBS9 in the densities of bovine BBSome **(D)**and human core BBSome **(E)**show that the hairpin reorients towards BBS8 in the absence of BBS2 and BBS7. **(F)**: Fitting the crystal structure of the BBS1 β-propeller in complex with Arl6 23 into the corresponding density of the BBS1 β-propeller in bovine BBSome indicates a clash of the Arl6 binding site with BBS2 and BBS7 25. **(G)**: An overlay of the orientation of the BBS1 β-propeller in human core BBSome with the density of the bovine BBSome shows a ~20° change in orientation **(H)**, which is sufficient to avoid the clash of the Arl6 binding site with BBS2 and BBS7.

**Figure S7:**
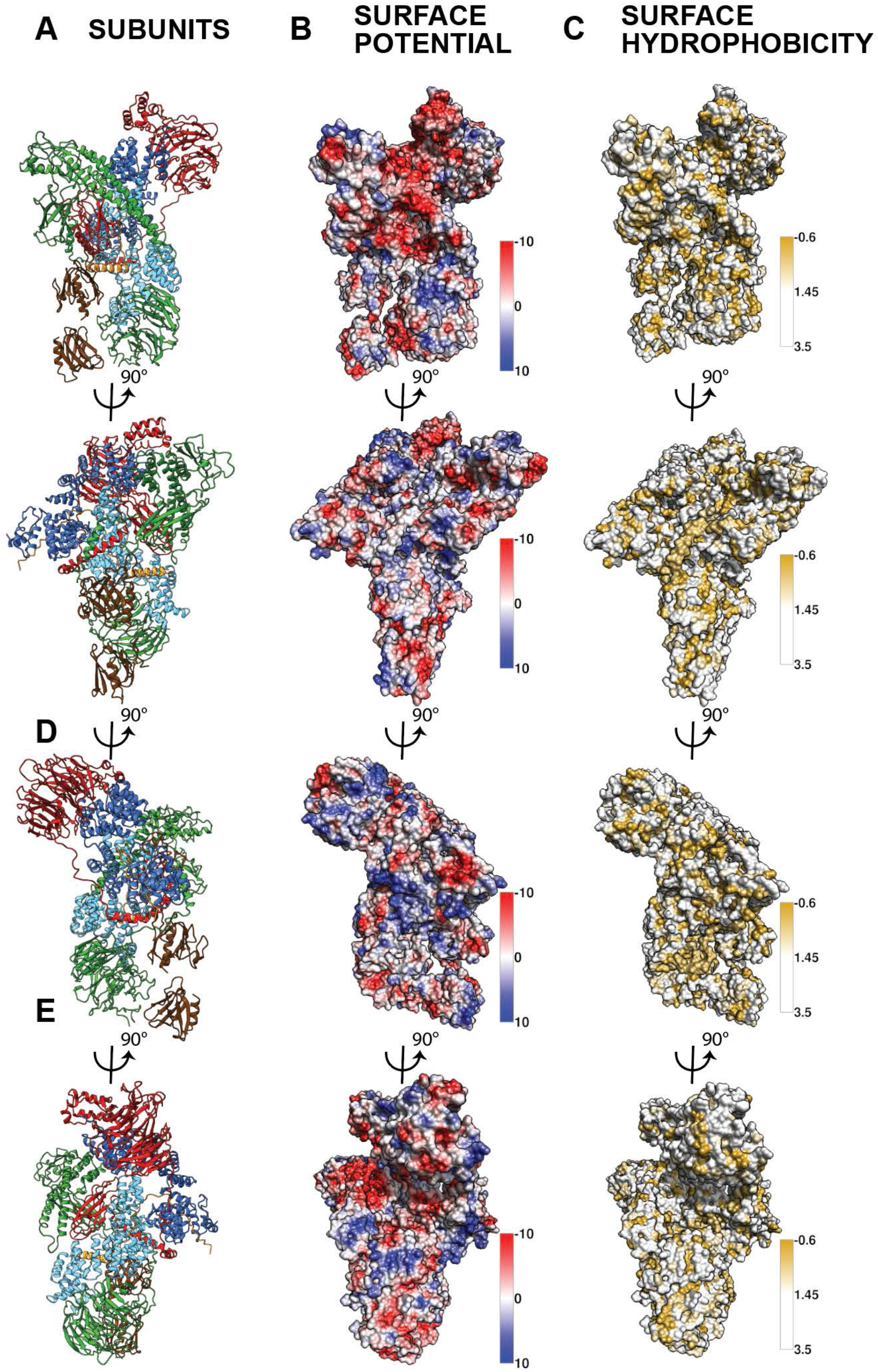
Surface potential and surface hydrophobicity of the core BBSome.

**Table S1:**
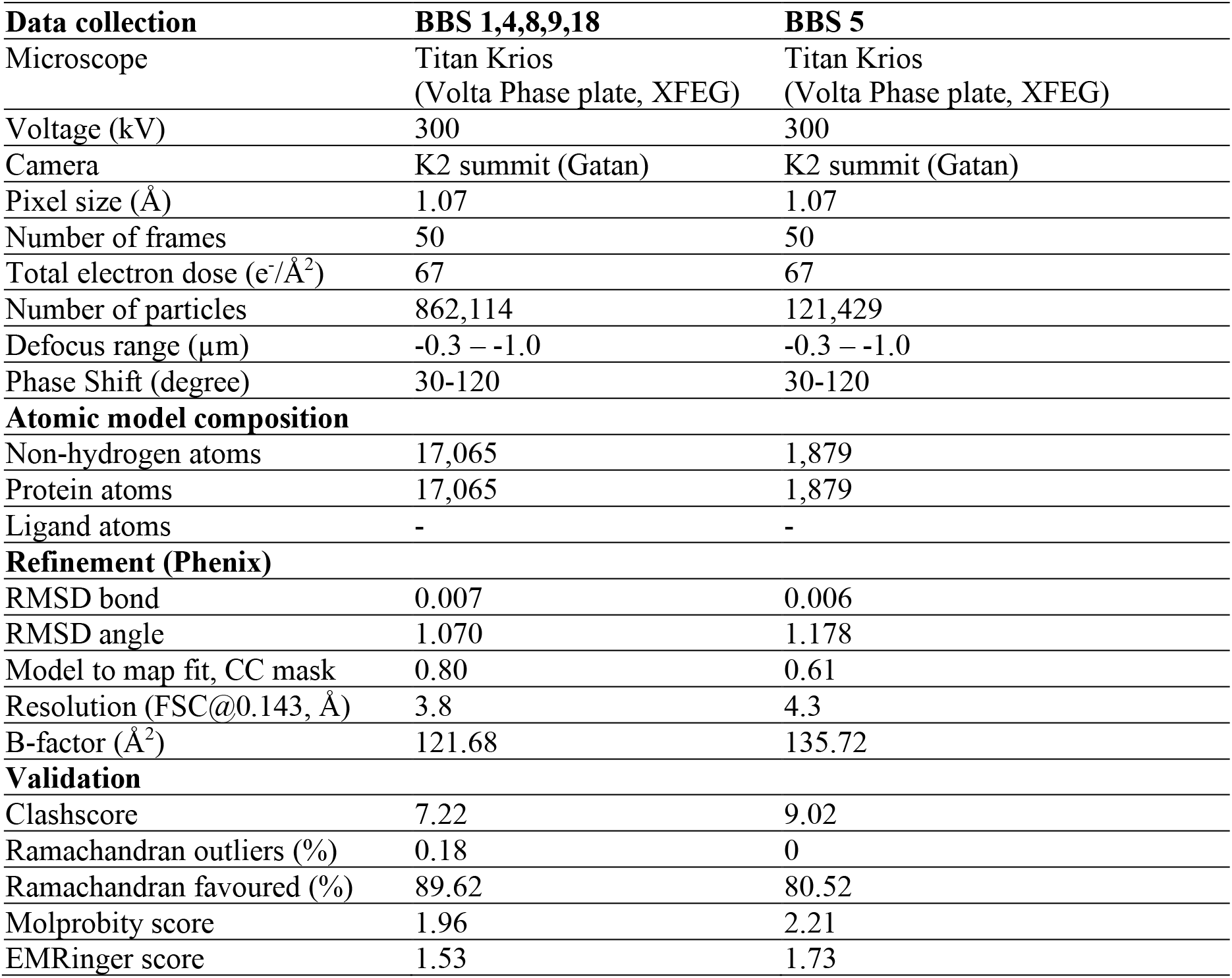
EM data collection and refinement statistics of the core BBSome. The BBS subunits 1,4,8,9 and 18 were modelled into the reconstruction from all particles that remained after ISAC 2D sorting, while the subunit BBS5 was modelled into a subset of particles derived by 3D clustering.

**Table S2:** Interfaces between core BBSome subunits. The mutual subunit interaction surfaces within the core BBSome with highest relevance for complex stability (i.e. with solvation free energies (Δ^i^G) < −2.5 kcal/mol and with interface areas > 400 Å^2^), as analyzed by the Pisa server 28.

| **BBSome**<br>**subunit A** | **BBSome**<br>**subunit B** | **Interface area**<br>**(A-B) [$\text{\AA}^2$]** | **$\Delta^iG$ (A-B)**<br>**[kcal/mol]** |
| --- | --- | --- | --- |
| BBS18 | BBS8 | 1621.0 | -28.1 |
| BBS18 | BBS4 | 1411.0 | -25.4 |
| BBS9 | BBS1 | 1946.2 | -23.0 |
| BBS9 | BBS8 | 2148.3 | -22.7 |
| BBS8 | BBS1 | 1785.8 | -20.5 |
| BBS4 | BBS1 | 2429.1 | -18.0 |
| BBS8 | BBS4 | 1469.4 | -13.1 |

